# T3SS effector VopL inhibits the host ROS response, promoting the intracellular survival of *Vibrio parahaemolyticus*

**DOI:** 10.1101/104315

**Authors:** Marcela de Souza Santos, Dor Salomon, Kim Orth

## Abstract

The production of antimicrobial reactive oxygen species by the nicotinamide dinucleotide phosphate (NADPH) oxidase complex is an important mechanism for control of invading pathogens. Herein, we show that the gastrointestinal pathogen *Vibrio parahaemolyticus* counteracts reactive oxygen species (ROS) production using the Type III Secretion System 2 (T3SS2) effector VopL. In the absence of VopL, intracellular *V. parahaemolyticus* undergo ROS-dependent filamentation, with concurrent limited growth. During infection, VopL assembles actin into non-functional filaments resulting in a dysfunctional actin cytoskeleton that can no longer mediate the assembly of the NADPH oxidase at the cell membrane, thereby limiting ROS production. This is the first example of how a T3SS2 effector contributes to the intracellular survival of *V. parahaemolyticus* to support the establishment of a protective intracellular replicative niche.

## Introduction

*Vibrio parahaemolyticus* is a Gram-negative bacterium that inhabits warm marine and estuarine environments throughout the world (1). This bacterium is recognized as the world’s leading cause of acute gastroenteritis associated with the consumption of contaminated raw or undercooked seafood (2). In immunocompetent individuals, the illness is self-limiting with symptoms including diarrhea with abdominal cramping, nausea, vomiting, and low-grade fever (1). However, for individuals with underlying health conditions, the bacterium can breach the gut barrier and cause septicemia corresponding to high mortality rates (3). *V. parahaemolyticus* has also been reported to cause infection of seawater-exposed wounds, which in rare cases escalates to necrotizing fasciitis and septicemia (4). The bacterium was also identified as the etiologic agent of acute hepatopancreatic necrosis disease (AHPND), a shrimp illness that has recently emerged, causing a massive economic burden on the shrimp industry (5).

Among several virulence factors, including thermostable hemolysins (TDH/TRH), polar and lateral flagella, and adhesins, *V. parahaemolyticus* encodes two Type III Secretion Systems (T3SS1 and T3SS2) (6). These are needle-like apparatuses used by the bacterium to inject proteins, termed effectors, into the host cell (7). The first T3SS, T3SS1, is present in all sequenced *V. parahaemolyticus* strains, including both environmental and clinical isolates, and is induced by culturing the bacteria in low Ca^2+^, as in serum-free Dulbecco’s modified Eagle’s medium (DMEM) tissue culture growth medium (8). While this system does not contribute to the bacterium’s enterotoxicity (9), the T3SS1 effectors orchestrate a multifaceted and efficient death of the infected host cell (10). *V. parahaemolyticus* more recently acquired the second T3SS, T3SS2, through a lateral gene transfer event and this system is primarily associated with clinical isolates (6). The T3SS2 becomes activated in the presence of bile salts (11, 12) and is recognized as the principal virulence factor causing gastroenteritis (9).

We recently reported that during infection, T3SS2 promotes *V. parahaemolyticus* invasion of non-phagocytic cells (13, 14). We found that *V. parahaemolyticus* encodes VopC (VPA1321), a deamidase that constitutively activates the GTPases Rac and Cdc42 resulting in membrane ruffling and uptake of the bacterium into the cell (14, 15). Once inside the host cell, the bacterium is initially contained within an endosome-like vacuole (13). Upon acidification of the vacuole, but prior to endosome fusion with the lysosome, *V. parahaemolyticus* breaks out of its vacuole and escapes into the cytosol (13). *V. parahaemolyticus* then uses the cell as a protected replicative niche (100–300 bacteria/cell) (13). Although historically studied as an exclusive extracellular bacterium, these findings changed this long-standing view and established *V. parahaemolyticus* as a facultative intracellular bacterium. While the role of VopC to promote host cell invasion is well-defined (14), the contribution of other T3SS2 effectors to the maintenance of the intracellular lifecycle of *V. parahaemolyticus* remains poorly understood.

VopL (VPA1370), a T3SS2 effector, encodes three consecutive WASP-homology 2 (WH2) domains intermixed with three proline-rich regions and a subsequent VopL C-terminal domain (VCD) (Fig. 1A) (16–18). WH2 domains are commonly found in nucleators of actin filaments; indeed, VopL’s *in vitro* nucleating activity is even more potent than that of the maximally-activated Arp2/3 complex (16). Ectopic expression of VopL in epithelial cells causes a dramatic rearrangement of the actin cytoskeleton into filaments reminiscent of stress fibers (16). Recent studies on VopF, the *V. cholerae* homolog of VopL, revealed that VopL/F works to sequester actin monomers and arrange those monomers into linear strings of non-functional filaments, as opposed to *bona fide* actin filaments (19). Despite the comprehensive characterization of VopL from a structural and a mechanistic standpoint, the contribution of this effector during a *V. parahaemolyticus* infection remained elusive.

Herein, we show that VopL is required for the intracellular survival of *V. parahaemolyticus*. In the absence of VopL, intracellular bacteria filament, which indicates that the bacteria are under stress. We identified the stressor as an increase in exposure to reactive-oxygen species (ROS). We found that the presence of VopL prevented filamentation by suppressing the generation of ROS by the nicotinamide adenine dinucleotide phosphate (NADPH) oxidase complex. To generate ROS, cytosolic and membranous subunits of the NADPH oxidase complex must come together at cell membranes (20). VopL, by hijacking the actin cytoskeleton, impedes the translocation of NADPH cytosolic subunits to the cell membrane, thereby preventing the complete assembly of the enzymatic ROS complex. Thus, *V. parahaemolyticus* deploys the T3SS2 effector VopL to secure a safe replicative niche within the host cell.

## Results

### In the absence of VopL, *V. parahaemolyticus* undergoes filamentation with concurrent decreased intracellular survival

As discussed earlier, despite not contributing to *V. parahaemolyticus*’ enterotoxicity (9), the T3SS1 becomes activated upon suspension in DMEM, causing the rapid death (~3h post-infection) of tissue-cultured cells (10), thereby masking the activity of the T3SS2. To reveal the activity of T3SS2 we made use of the *V. parahaemolyticus* CAB2 strain, an isogenic strain derived from the clinical isolate RimD2210633 (14). CAB2 contains a deletion for genes encoding two types of toxic factors. First, a deletion was made in *exsA* loci, resulting in the inactivation of the transcriptional activator for the T3SS1 (21). Second, deletions were made for *tdhAS*, the two thermostable direct hemolysins (TDH) present in RimD2210633, eliminating their cytotoxic activity (22). The resulting strain, CAB2, has been used in subsequentstudies to assess the activity of T3SS2 and its effectors.

Initially, we set out to investigate the contribution of VopL for the survival of CAB2 within Caco-2 cells, a colonic epithelial cell line. Bacterial survival can be assessed in two ways: first by determining the number of intracellular bacteria as a function of time post-invasion (Fig. 1B) and second by visualization of intracellular CAB2 using confocal microscopy (Fig. 1C and Fig. S1A,B). Shortly after the invasion of Caco-2 cells (2h post-invasion), CAB2 and its VopL-mutant counterpart (CAB2Δ*vopL*) exhibited comparable cell invasion and intracellular growth (Fig. 1B), displayinga patchy distribution inside their host cell (Fig. S1A). This activity was reminiscent of *V. parahaemolyticus*’ vacuolar localization early after invasion (13). At 4h post-invasion, CAB2 and CAB2Δ*vopL* counts doubled (Fig. 1B), consistent with intracellular replication, which was indicated by a significant increase in bacterial load within Caco-2 cells (Fig. S1B). At 6h post-invasion, CAB2 exhibited an additional growth increase, while CAB2Δ*vopL* survival was substantially compromised as bacterial counts dropped more than twofold (Fig. 1B). Strikingly, confocal inspection of intracellular CAB2Δ*vopL* at 6h post-invasion revealed a dramatic change in bacterial morphology: CAB2 displayed characteristic *V. parahaemolyticus’* rod-shape, while CAB2Δ*vopL* appeared significantly elongated (Fig. 1C). Expression of VopL in CAB2Δ*vopL* from a plasmid (CAB2Δ*vopL*+pVopL) increased intracellular growth to levels comparable to that of wild type bacteria (Fig. 1B) and also rescued normal bacterial morphology (Figs. 1C,D).

Elongation of CAB2Δ*vopL* was not a result of the cell body extension, but rather a deficiency in bacterial cell division. Closer inspection of CAB2Δ*vopL* revealed that the elongated cell body contained multiple nucleoids (Fig. S1C), which is consistent with continuous bacterial replication but ceased septation. This morphological phenotype is referred to as bacterial filamentation and represents an important strategy used by bacteria to survive during stressful situations (23). Several bacteria have been reported to undergo filamentation as a protective mechanism against phagocytosis, as in the case of Uropathogenic *Escherichia coli* (24), or against consumption of “inedible” filamentous bacteria by protists, as in the case of *Flectobacillus* spp. (25). Filamentation can also be triggered in response to DNA-damaging stresses such as UV radiation, antibiotics, and reactive oxygen species (ROS) (23). Given that CAB2Δ*vopL* deliberately invades Caco-2 cells (via VopC), samples are not exposed to UV radiation, and intracellular bacteria are not exposed to gentamicin, as this antibiotic is not taken up by Caco-2 cells, we hypothesized that host generation of ROS could be responsible for the filamentous CAB2Δ*vopL*.

**Figure 1.**
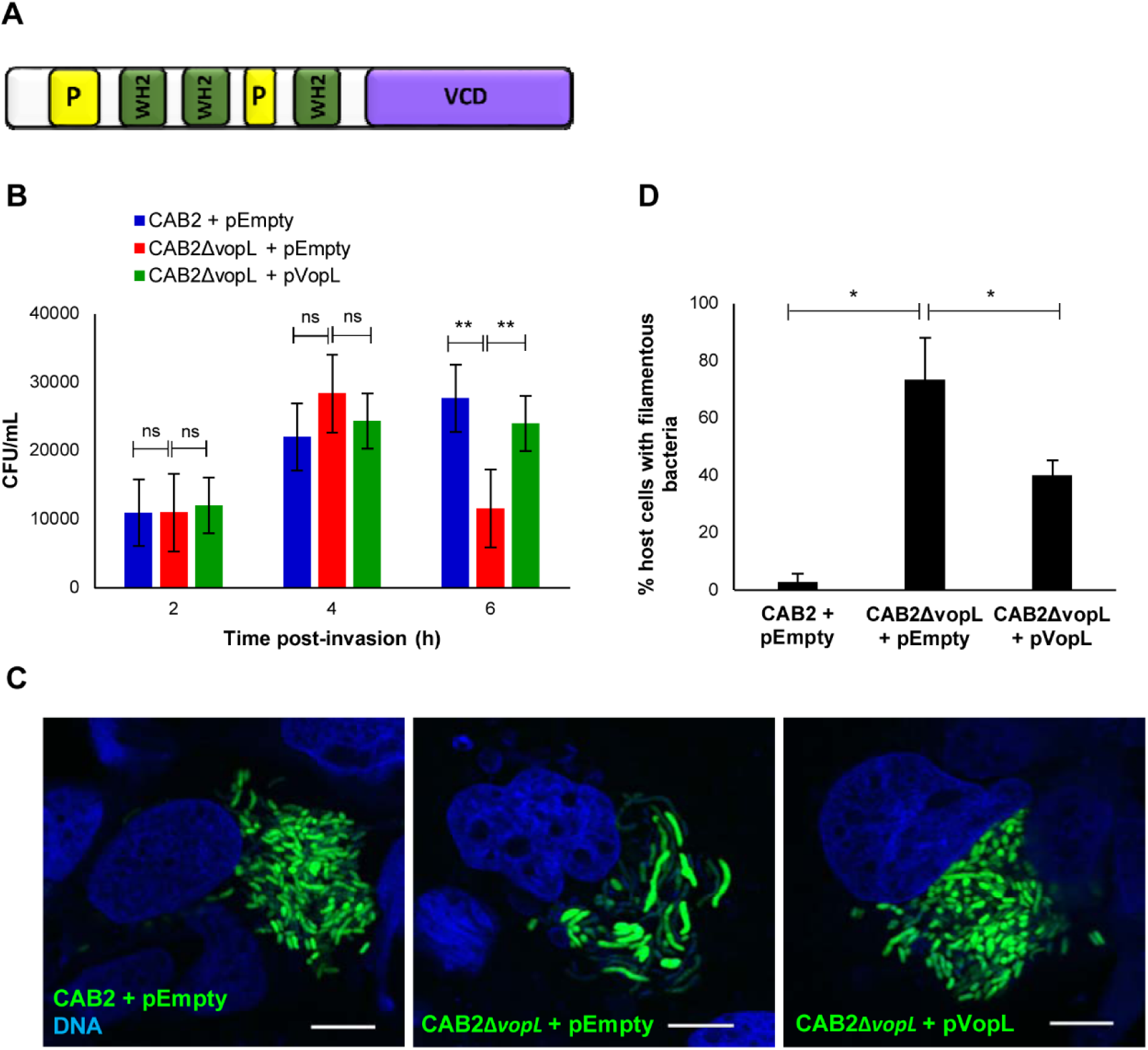
Intracellular CAB2 undergoes filamentation with concurrent decreased survival in the absence of VopL. **(A)** Schematic of VopL domains: WASP-homology 2 domains (WH2), proline-rich regions (P), and VopL C-terminal domain (VCD). **(B)** Intracellular growth of CAB2 strains in Caco-2 cells. Caco-2 cells were infected with indicated CAB2 strains for 2h followed by incubation with 100 μg/mL gentamicin for the specified times. Cell lysates were serially diluted and plated and CFU was enumerated for intracellular bacteria. Values are means ± SD of a representative experiment performed in triplicate. Asterisks indicate statistically significant difference in intracellular bacteria CFU counts between CAB2 + pEmpty and CAB2Δ*vopL* + pEmpty or between CAB2Δ*vopL* + pEmpty and CAB2Δ*vopL* + pVopL. ** *p* < 0.005. **(C)** Confocal micrographs of Caco-2 cells infected with indicated GFP-tagged CAB2 strains for 2h and incubated with 100 μg/mL gentamicin for 6h. DNA was stained with Hoechst (blue). Scale bars, 10 μm. **(D)** Caco-2 cells invaded by CAB2 + pEmpty, CAB2Δ*vopL* + pEmpty, or CAB2Δ*vopL* + pVopL were enumerated for containment of filamentous bacteria. 90 cells per sample (each sample referring to infection by one of the 3 bacterial strains) were enumerated over 3 independent experiments. Values are means ± SD. Asterisks indicate statistically significant difference between CAB2- and CAB2Δ*vopL*-infection samples (* *p* = 0.0116) or between CAB2Δ*vopL*- and CAB2Δ*vopL* + pVopL-infection samples (* *p* = 0.0421).

### Bacterial filamentation is ROS-dependent

ROS is produced by NADPH oxidases, specialized enzymes whose sole function is the generation of ROS (20). There are seven members of the NADPH oxidase (NOX) family, NOX1–5 and two dual oxidases (DUOX1 and DUOX2), which collectively produce ROS in a wide range of tissues where ROS participate in a variety of cell processes such as mitogenesis, apoptosis, hormone synthesis, and oxygen sensing (20, 26). NOX2 is a phagocyte-specific isoform, being highly expressed in neutrophils and macrophages where it plays an essential role in host defense against microbial pathogens (20). NOX1 is the closest homolog of NOX2, with whom it shares 56% sequence identity (20). NOX1 is most abundant in the colon epithelium and is also expressed in a variety of cell lines, including Caco-2 cells (26, 27). At present, the physiological roles of colonic NOX1 are not fully understood. NOX1-derived ROS has been implicated in control of cell proliferation, mucosal repair after injury, and inflammatory response (26). Importantly, evidence suggests a role for NOX1 as a host defense oxidase (28). For instance, colon epithelial cells exhibited high NOX1-mediated ROS production in response to flagellin from *Salmonella enteriditis* (29).

To assess whether filamentation of CAB2Δ*vopL* within Caco-2 cells was resultant from bacterial exposure to ROS, we suppressed ROS generation using GKT136901 (GKT), a direct and specific inhibitor of NOX1/4 (NOX4 is primarily expressed in the kidney (20)) (30). Caco-2 treatment with 10 μM GKT reduced the number of host cells containing filamentous CAB2Δ*vopL* by more than twofold (Fig. 2), suggesting that NOX1-generated ROS mediates bacterial filamentation.

**Figure 2.**
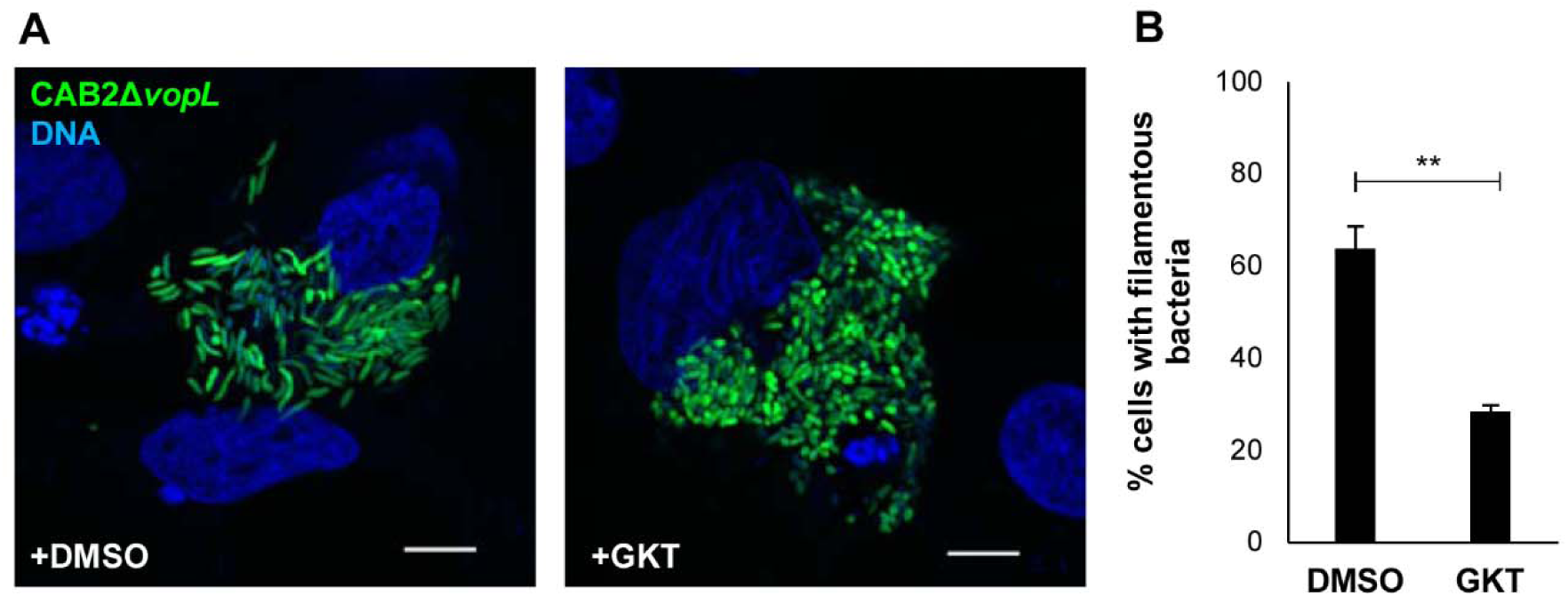
CAB2Δ*vopL* filamentation is ROS-dependent. **(A)** Confocal micrographs of Caco-2 cells infected with CAB2Δ*vopL*-GFP (green) for 2h followed by incubation with 100 μg/mL gentamicin for 6h. Host cells were pre-treated with either dimethyl sulfoxide (DMSO) or 10 μM GKT, which were kept throughout infection. DNA was stained with Hoechst (blue). Scale bars, 10 μm. (**B**) Quantification of filamentous bacteria in the presence or absence of GKT. Caco-2 cells invaded by CAB2Δ*vopL*-GFP and treated with either DMSO or GKT were analyzed for presence of filamentous bacteria. 300 cells for each sample (DMSO or GKT), over 3 independent experiments, were analyzed for presence of filamentous bacteria. Values are means ± SD. Asterisks indicate statistically significant difference between DMSO and GKT samples (** *p* = 0.0038).

### VopL is required for normal bacterial growth in COS^phox^ cells

Our results thus far show that NOX1-generated ROS is causal for bacterial filamentation and that filamentation only occurs in the absence of VopL. Therefore, we hypothesized that VopL suppresses generation of ROS. To quantify NADPH oxidase-dependent production of ROS, we analyzed host cell release of superoxide, the product of NADPH oxidase-mediated reduction of molecular oxygen and the precursor of other ROS (31). Detection of ROS generated by endogenous NOX1 in the colon, as well as in Caco-2 cells, was challenging to measure (29). Indeed, under our experimental conditions we could not detect superoxide in a sensitive manner during infection of Caco-2 cells, nor could we detect it upon cell stimulation with the PKC activatior phorbol 12-myristate 13-acetate (PMA) (Fig. S2).

Thus, to investigate the ability of VopL to control the generation of ROS by NADPH oxidases, we used a well characterized model cell system, the COS^phox^ cell line, that has been used previously to biochemically analyze the production of ROS (32). These cells stably express NOX2 (gp91^phox^) along with the other NOX2 complex subunits p22^phox^, p47^phox^, and p67^phox^ (32). Notably, the NOX1 enzymatic complex also includes p22^phox^, along with NOXO1 and NOXA1, homologs of p47^phox^ and p67^phox^, respectively (27, 33). Given the similarity in functioning of the NOX1 and NOX2 complexes, COS^phox^ cells represent a suitable model of non-phagocytic cells with robust NOX-dependent production of ROS for the present study.

Initially, we investigated CAB2 growth within COS^phox^ cells, in the presence and absence of VopL. While CAB2 was able to efficiently replicate inside COS^phox^ cells (Fig. 3A) and displayed the bacterium’s normal rod-shape (Fig. 3B and Fig. S3), CAB2Δ*vopL* grew at approximately half the rate of wild type bacteria (Fig. 3A) and, importantly, demonstrated a very dramatic filamentous phenotype (Fig. 3B and Fig. S3). Expression of VopL in CAB2Δ*vopL* from a plasmid (CAB2Δ*vopL +* pVopL) rescued intracellular growth to levels comparable to that of wild type bacteria (Fig. 3A) and restored the rod-shaped bacterial morphology (Fig. 3B,C). These findings are in agreement with the observations made using Caco-2 cells and support a role for VopL in bacterial intracellular survival.

**Figure 3.**
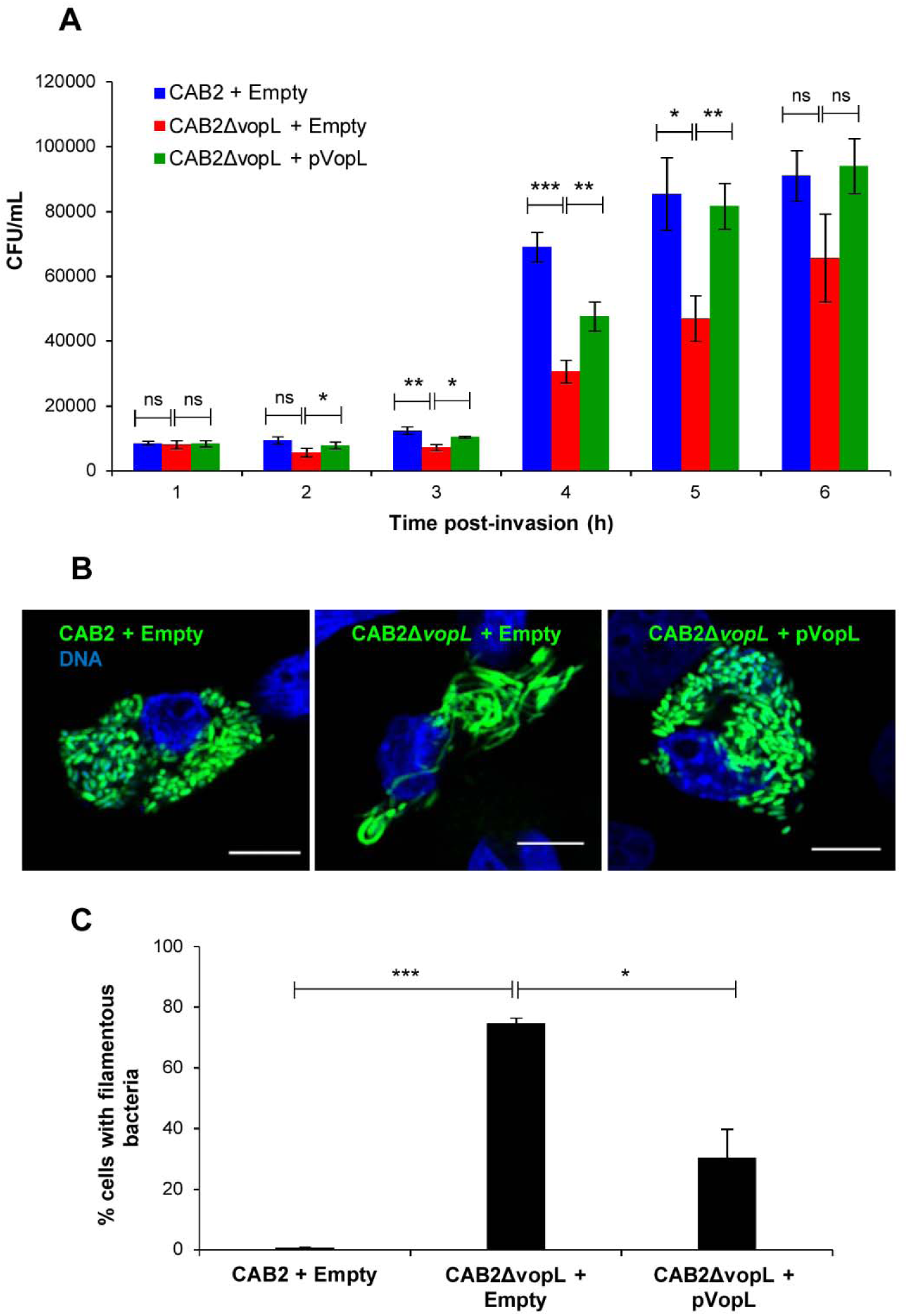
VopL is required for CAB2 survival within COS^phox^ cells. **(A)** Intracellular growth of CAB2 strains in COS^phox^ cells. COS^phox^ cells were infected with indicated CAB2 strains for 2h followed by incubation with 100 μg/mL gentamicin for the specified times. Cell lysates were serially diluted and plated and CFU was enumerated for intracellular bacteria. Values are means ± SD of a representative experiment performed in triplicate. Asterisks indicate statistically significant difference in intracellular bacteria CFU counts between CAB2 + pEmpty and CAB2Δ*vopL* + pEmpty or between CAB2Δ*vopL* + pEmpty and CAB2Δ*vopL* + pVopL. * *p* < 0.05, ** *p* < 0.005, *** *p* < 0.0005. **(B)** Confocal micrographs of COS^phox^ cells infected with indicated GFP-tagged CAB2 strains for 2h and incubated with 100 μg/mL gentamicin for 6h. DNA was stained with Hoechst (blue). Scale bars, 10 μm. **(C)** COS^phox^ cells invaded by CAB2 + pEmpty, CAB2Δ*vopL* + pEmpty, or CAB2Δ*vopL* + pVopL were enumerated for containment of filamentous bacteria. 90 cells per sample (each sample referring to infection by one of the 3 bacterial strains) were enumerated over 3 independent experiments. Values are means ± SD. Asterisks indicate statistically significant difference between CAB2- and CAB2Δ*vopL*-infection samples (*** *p* = 0.0002) or between CAB2Δ*vopL*- and CAB2Δ*vopL +* pVopL-infection samples (* *p* = 0.0122).

### Filamentation of Δ*vopL V. parahaemolyticus* within COS^phox^ cells is ROS-dependent

As with Caco-2 cells, we investigated whether filamentation of intracellular CAB2Δ*vopL* resulted from ROS generation by COS^phox^ cells. In the case of COS^phox^ cells, NOX2-dependent generation of ROS was inhibited by apocynin (APO), which blocks ROS generation by preventing the complete assembly of the NOX2 enzymatic complex (20). APO treatment of COS^phox^ cells abrogated infection-elicited generation of superoxide (Fig. S4) and significantly attenuated CAB2Δ*vopL* filamentation (Fig. 4A), reducing the number of COS^phox^ cells containing filamentous bacteria by 40% (Fig. 4B). Therefore, in agreement with our findings obtained with Caco-2 cell infection, ROS is an agent involved bacterial filamentation.

### VopL suppresses generation of NOX2-derived superoxide

As a control for NOX2-dependent generation of superoxide, we stimulated COS^phox^ cells with PMA. PMA activates the NOX2 complex via PKC-mediated phosphorylation of the p47^phox^ subunit (34). PMA stimulation significantly enhanced superoxide production in comparison to untreated cells (Fig. 4C). Infection of COS^phox^ with CAB2 induced generation of superoxide at levels approximately 1.5 times higher than uninfected cells (Fig. 4C). Strikingly, cells infected with CAB2Δ*vopL* released 3 times more superoxide than uninfected cells (Fig. 4C). Rescue of the CAB2Δ*vopL* strain with a VopL-expression plasmid lowered superoxide production to levels similar to that generated by the parental CAB2 strain (Fig. 4C). These results confirm our hypothesis that the presence of VopL suppresses NOX-generated ROS.

**Figure 4.**
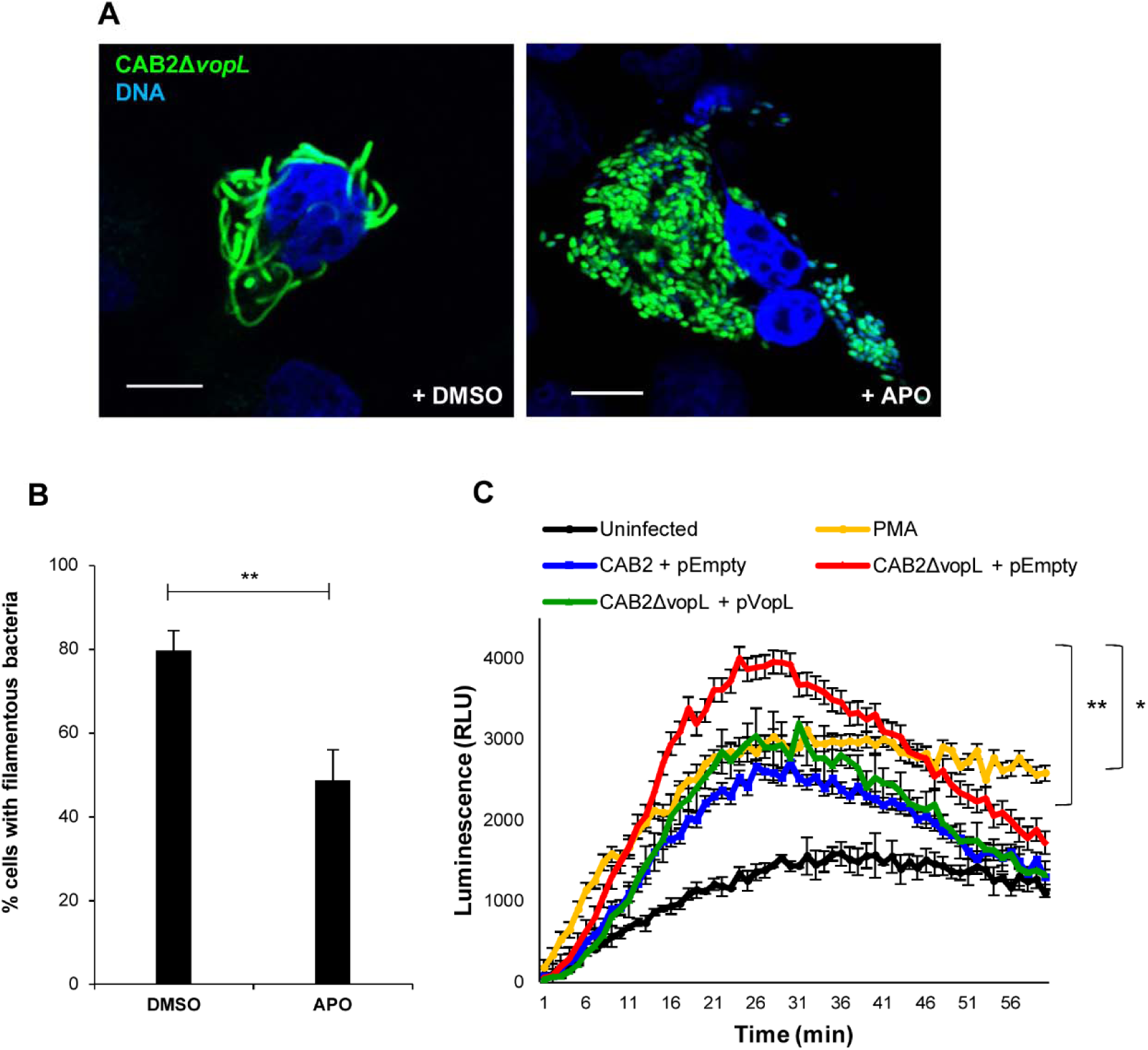
VopL inhibits generation of superoxide. **(A)** Confocal micrographs of COS^phox^ cells infected with CAB2Δ*vopL*-GFP (green) for 2h followed by incubation with 100 μg/mL gentamicin for 6h. At the beginning of the last hour of gentamicin incubation, either dimethyl sulfoxide (DMSO) or 250 μM APO was added to the samples. DNA was stained with Hoechst (blue). Scale bars, 10 μm. **(B)** Quantification of filamentous bacteria in the presence or absence of APO. COS^phox^ cells invaded by CAB2Δ*vopL*-GFP and treated with either DMSO or APO were analyzed for presence of filamentous bacteria. 300 cells for each sample (DMSO or APO), over 3 independent experiments, were analyzed for presence of filamentous bacteria. Values are means ± SD. Asterisks indicate statistically significant difference between DMSO and APO samples (** *p* = 0.0058). **(C)** COS^phox^ cells were infected with indicated CAB2 strains for 2h and superoxide production was measured as a function of luminescence intensity. Values are means ± SD from one representative experiment. Asterisks indicate statistically significant difference between CAB2- and CAB2Δ*vopL*-infection samples (** *p* = 0.0039) or between CAB2Δ*vopL*- and CAB2Δ*vopL +* pVopL-infection samples (* *p* = 0.0387).

### VopL suppresses the movement of NOX2 cytosolic subunits to cell membranes

As mentioned earlier, NOX2 is a multi-subunit complex; the latent complex is disassembled in resting cells and must become assembled at cell membranes with potential to generate ROS (20). When at rest, the NOX2 complex regulatory subunits p67^phox^ and p47^phox^ are present in the cytosol as a heterotrimeric complex along with p40^phox^ (20). Rac, another complex subunit, is also present in the cytosol in its inactive, GDP-bound, form. Upon cell stimulation, all activated cytosolic components translocate to both cell plasma and phagocytic membranes where they interact with membranous subunits gp91^phox^ (NOX2) and p22^phox^ and complete the assembly of a functional NOX complex (20). Several pieces of evidence support a role for actin in NOX activity. For instance, addition of G-actin was shown to potentiate NOX activity in a cell-free system (35). The p47^phox^ and p67^phox^ each contain a SH3 domain known to associate with the actin cytoskeleton (34). In resting polymorphonuclear leukocytes (PMN), p67^phox^ is detected exclusively in the detergent-insoluble, cytoskeletal fraction (34). Additionally, inhibition of actin polymerization by cytochalasin has been shown to modulate the translocation of NOX2 subunits from the cytosol to the plasma membrane upon stimulation of PMNs (36, 37).

Given that VopL disrupts the normal assembly of the actin cytoskeleton (19), we hypothesized that VopL would impair the complete assembly of the NOX2 complex, therefore compromising the production of ROS. To test this hypothesis, we transiently transfected COS^phox^ cells with VopL and subsequently induced NOX2 activation using PMA. Cell stimulation with PMA caused p67^phox^ to translocate from the cytosol to the plasma membrane and also caused an extensive rearrangement of the actin cytoskeleton with formation of membrane ruffles that co-localized with p67^phox^ (compare Figs. S5A and S5B). As previously reported (16), ectopic expression of wild type VopL (WT VopL) caused the formation of long actin strings reminiscent of stress fibers (Fig. 5A and Fig. S5C). In cells transfected with WT VopL, PMA-stimulated actin ruffles were not formed (Fig. 5A) and, importantly, the translocation of p67^phox^ from the cytosol to the cell membrane was impaired (Fig. 5A). Quantification of enrichment of p67^phox^ at the plasma membrane revealed a significantly smaller presence of this subunit at the membrane in the presence of VopL (Fig. 5B).

To confirm that the inhibitory effect of VopL on p67^phox^ translocation is dependent on the effector’s ability to manipulate the actin cytoskeleton, we also transfected cells with WH2*3-VopL, which contains point mutations at amino acids required for the actin binding activity of the WH2 domains (16). We previously established that WH2*3-VopL is devoid of actin assembly activity *in vitro* and does not induce actin stress fiber formation in transfected cells (16) (Fig. S5D). Expression of WH2*3-VopL impaired neither PMA-stimulated membrane ruffling nor the cytosol-plasma membrane translocation of p67^phox^ (Fig. 5C), which was highly enriched at the plasma membrane (Fig. 5D).

Therefore, VopL, by disrupting the normal assembly of the actin cytoskeleton, impairs the assembly of the NOX2 complex, thereby limiting the generation of ROS induced during infection.

**Figure 5.**
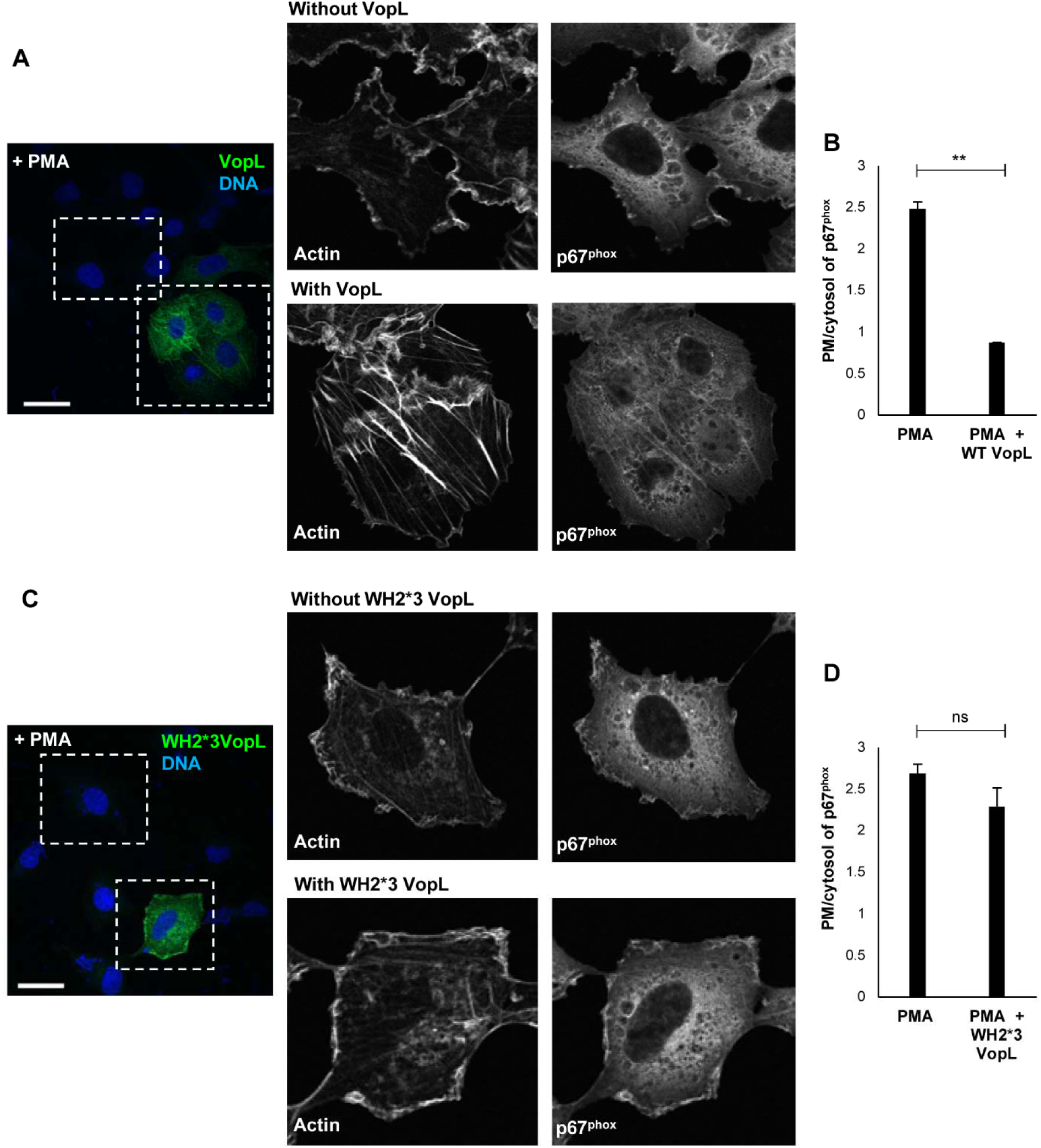
VopL inhibits assembly of NOX2 complex. COS^phox^ cells were transiently transfected with either wild type VopL (WT VopL, panel **A**) or catalytically inactive VopL (WH2*3-VopL, panel **C**) and subsequently stimulated for ROS response with 0.4 μg/mL phorbol 12-myristate 13-acetate (PMA). Cells were immunostained for p67^phox^ (colored in gray scale to enhance contrast) and VopL (green). DNA and actin were stained with Hoechst (blue) and Alexa Fluor 680 phalloidin (colored in gray scale to enhance contrast), respectively. White, dotted boxes highlight regions within untransfected and transfected cells that were magnified. Scale bars, 40 μm. **(B,D)** PMA-stimulated translocation of p67^phox^ from the cytosol to the plasma membrane, for untransfected cells and cells transfected with either WT VopL **(B)** or WH2*3-VopL **(D)**, was quantified by analysis of line scans crossing the two cellular compartments. 90 cells for each population (untransfected or transfected with VopL) were analyzed over 3 independent experiments. Values are means ± SD. Asterisk indicates statistically significant difference between untransfected and WT VopL-transfected cells (** *p*=0.0011).

### Jasplakinolide phenocopies VopL-mediated inhibition of NOX2 complex assembly

VopL rearranges the actin cytoskeleton into linear strings of non-functional filaments that resemble stress fibers (Fig. 5A). By doing so, VopL retains p67^phox^ in the cytosol, and consequently, impedes the activation of NOX2. Therefore, we assessed whether manipulation of the actin cytoskeleton in the form of stable stress fiber-like structures could solely account for the inhibition NOX2 assembly.

Jasplakinolide is a potent inducer of actin polymerization and a stabilizer of actin filaments (38). Treatment of COS^phox^ cells with jasplakinolide induced stress fiber formation (compare Figs. 6A and 6B). As with VopL, jasplakinolide treatment abrogated PMA-stimulated translocation of p67^phox^ from the cytosol to the plasma membrane (compare Figs. 6C and 6D). These findings further support our hypothesis that VopL manipulates the actin cytoskeleton to prevent NOX assembly and dampen ROS production.

**Figure 6.**
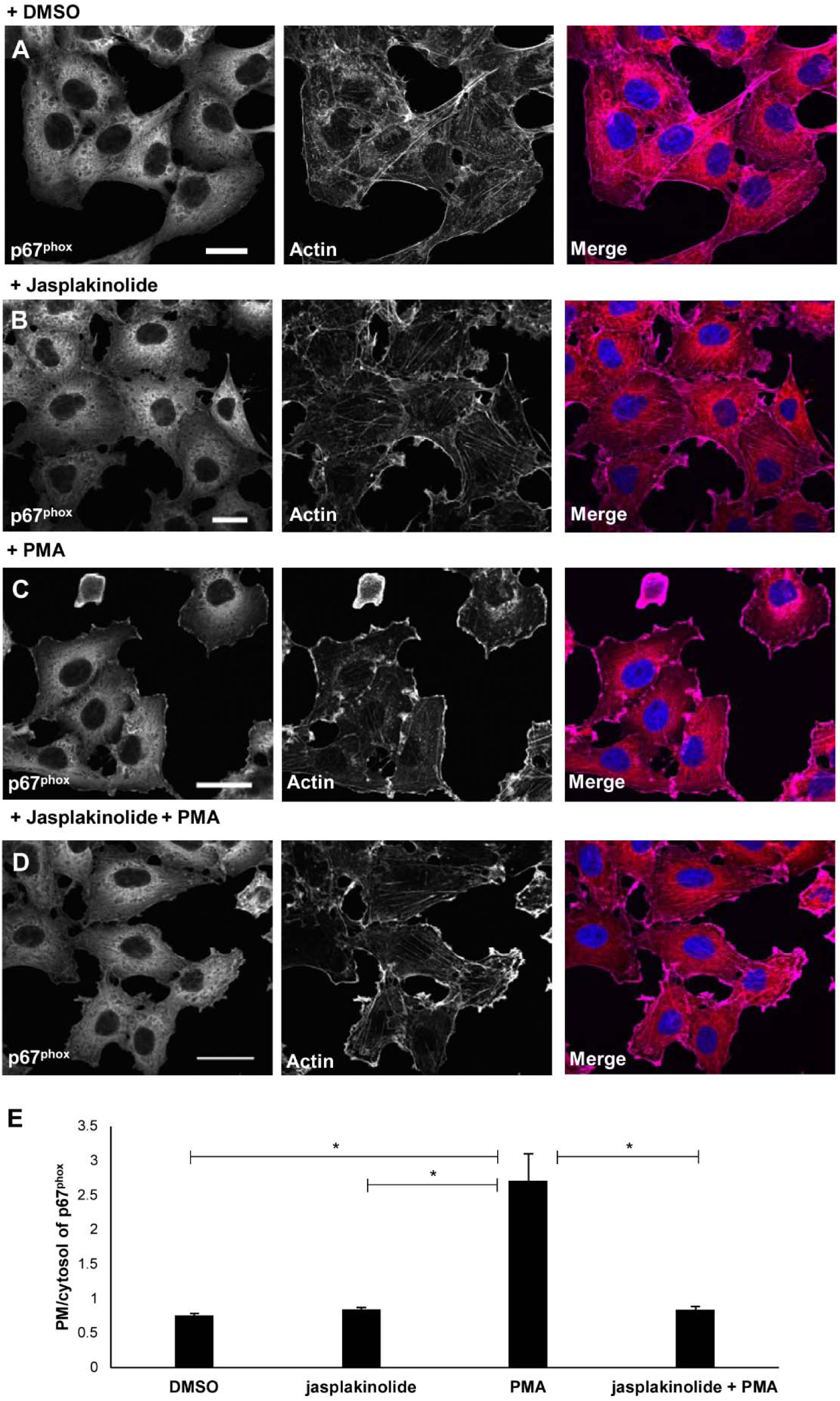
Jasplakinolide arrests assembly of NOX2 complex. COS^phox^ cells were treated with either DMSO **(A)** or 100 nM jasplakinolide **(B)** and subsequently stimulated for ROS response with 0.4 μg/mL phorbol 12-myristate 13-acetate (PMA) **(C,D)**. Cells were immunostained for p67^phox^ (colored in gray scale to enhance contrast). DNA and actin were stained with Hoechst (blue) and Alexa Fluor 680 phalloidin (colored in gray scale to enhance contrast), respectively. Scale bars, 40 μm. **(E)** PMA-stimulated translocation of p67^phox^ from the cytosol to the plasma membrane, for cells transfected with jasplakinolide or left untreated, was quantified by analysis of line scans crossing the two cellular compartments. 90 cells for each population (DMSO treatment only, jasplakinolide-treated only, PMA-stimulated, or jasplakinolide+PMA treated) were analyzed over 3 independent experiments. Values are means ± SD. Asterisk indicates statistically significant difference between mock (DMSO) or PMA stimulated cells (* *p*=0.0126), between jasplakinolide or PMA-treated cells (* *p*=0.0136), and between PMA-stimulated or jasplakinolide+PMA treated cells (* *p*=0.0127).

## Discussion

The T3SS2 effector VopL and its *V. cholera* homologue VopF were discovered about a decade ago and their structure and biochemical activities are now well-characterized (16–19, 39–41). The coincidental discoveries that VopL/F produces linear strings of non-functional filaments by Avvaru and colleagues (19) and our findings that *V. parahaemolyticus* is a facultative intracellular pathogen (13), laid the groundwork for the elucidation of the biological role of VopL during infection. Here, we showed that NOX-derived generation of ROS plays an important role in controlling intracellular proliferation of *V. parahaemolyticus*. Specifically, ROS-dependent stress of the bacterium resulted in impairment of cell division and consequent filamentation of the bacteria. VopL is essential in preventing this deleterious event: by directly targeting the actin cytoskeleton and catalyzing the assembly of non-canonical actin filaments, VopL arrests the actin-dependent movement of cytosolic NOX subunits to cell membranes. As a result, VopL prevents the activation of the NOX complex and consequent production of ROS. Thus, this is the first report of how VopL aids *V. parahaemolyticus* infection: it secures a relatively “stress-free” environment within the host cell, enabling the bacterium to establish a successful replicative niche.

Alternative methods are used by other intracellular pathogens to inhibit ROS-mediated killing. Some pathogens scavenge ROS using extracellular polysaccharides, as in the case of *Burkholderia cenocepacia* and *Pseudomonas aeruginosa* (42, 43). Several other pathogens act on signaling machinery upstream of ROS and prevent activation of the NOX complex. (44–47). *Salmonella enterica* Typhimurium excludes the NOX2 membranous subunits from the vacuole it inhabits in a T3SS-dependent manner (48). Importantly, the virulence factors and mechanisms used by the vast majority of these pathogens remain unknown. The present work not only identified the virulence factor used by *V. parahaemolyticus* to suppress host ROS generation, but also revealed an unprecedented mechanism used by a microbial pathogen to do so (Fig. 7).

The pharmacological inhibition of NOX1 and NOX2 complexes, shown to fully suppress ROS generation in the case of apocynin, did not result in complete reduction of bacterial filamentation. These findings raise the possibility that additional factors may contribute to the arrest of bacterial division. Intracellular growth of certain *Salmonella* strains resulted in filamentous bacteria due to a defect in the bacterial histidine biosynthetic pathway (49). Because *V. parahaemolyticus* only undergoes filamentation in the absence of VopL, under our experimental conditions, it is likely that host factors, rather than bacterial ones, elicit this morphological phenotype. Rosenberger and Finlay (50) reported an upregulation of MEK1 kinase during *S. enterica* Typhimurium infection of RAW 264.7 macrophages. MEK upregulation was causal for *Salmonella* filamentation, as occurrence of filamentous bacteria partially decreased in the presence of MEK inhibitors (50). MEK and NOX activities operated in parallel to mediate *Salmonella* filamentation (50). It is known that the actin cytoskeleton functions as a scaffold that mechanically modulates the activation of signaling pathways (51). For instance, pharmacological inhibition of the actin polymerization reportedly inhibited ERK and AKT activation (52). Interestingly, *V. parahaemolyticus* expresses VopA, a homolog of *Yersinia* spp. YopJ Ser/Thr acetyltransferase that has been shown to inhibit MAPK signaling pathways during infection (53–55). Therefore, in addition to NOX assembly and MAPK signaling pathways, other pathways will be the subject of future investigation as further mediators of filamentation of the VopL-mutant.

It is important to note that, despite the close homology between the NOX1 and NOX2 complexes (the closest homologs within the NOX family), NOXO1, the homolog of p47^phox^ in the NOX1 complex, lacks the autoinhibitory region (AIR) domain present in p47^phox^ (27, 33). As a result, NOXO1, as well as its partner NOXA1 (p67^phox^ homolog), are constitutively associated with p22^phox^ at the cell membrane (56). This accounts for the constitutive, stimulus-independent generation of ROS observed in the colon (27). Importantly, it has been shown that constitutive, NOX1-derived generation of ROS is partial and can be further enhanced upon PMA-stimulation of intestinal epithelial cells (29). In agreement with these findings, ectopic expression of NOX1, NOXO1, and NOXA1 across a range of cell types only results in partial, sometimes absent, production of ROS, requiring PMA-stimulation for maximal generation of ROS (27, 33). These findings indicate that although constitutive NOX1, p22phox, NOXO1, and NOXA1 complex formation may be sufficient for generation of ROS, stimulated activation (GDP to GTP switch) of Rac is important to optimize this process (57). For instance, PMA facilitates the formation of GTP-bound Rac (58). These findings support our model that VopL interferes with ROS production in Caco-2 by impeding GTP-Rac mediated, optimal activation of the NOX1 complex.

Over many years of evolution, *V. parahaemolyticus*’ T3SS2 has maintained a repertoire of around a dozen effectors. One of these, VopC, has been shown to mediate cell invasion, and we now show a role for VopL in relieving free-radical stress for this intracellular pathogen. The activities of some of the other T3SS2 effectors have been studied and now the role that they play in this evolutionarily conserved invasive T3SS are ripe for future investigation.

**Figure 7.**
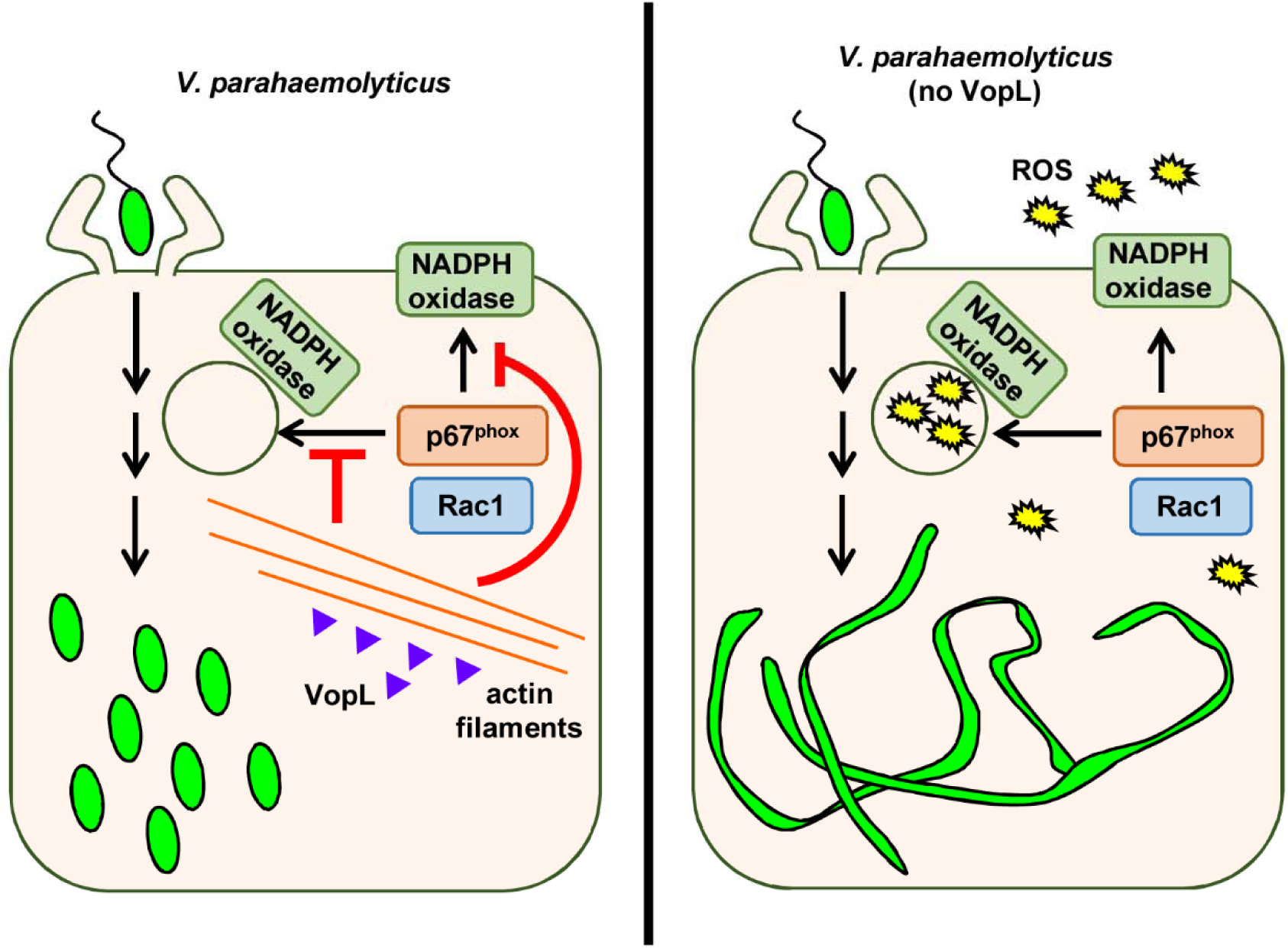
VopL inhibits NOX-derived production of ROS. *V. parahaemolyticus* deploys its T3SS2 effector VopL to disrupt the actin cytoskeleton. As a result, NOX cytosolic regulatory subunits do not translocate to cell membranes. In the absence of VopL, NOX-mediated generation of ROS induces bacterial stress, leading to bacterial filamentation.

## Materials and methods

### Bacterial strains and culture conditions

The *V. parahaemolyticus* CAB2 strain was derived from POR1 (clinical isolate RIMD2210633 lacking TDH toxins), the latter being a generous gift from Drs. Tetsuya Iida and Takeshi Honda (59). The CAB2 strain was made by deleting the gene encoding the transcriptional factor ExsA, which regulates the activation of the T3SS1 (14). CAB2 was grown in Luria-Bertani (LB) medium, supplemented with NaCl to a final concentration of 3% (w/v), at 30 °C. When necessary, the medium was supplemented with 50 μg/mL spectinomycin (to select for growth of CAB2-GFP strains (9)) or 250 μg/mL kanamycin.

### Deletion of *vopL* from CAB2 strain

For in-frame deletion of *vopL* (*vpa1370* in RimD2210633, GeneBank sequence accession number NC_004605), the nucleotide sequences 1kb upstream and downstream of the gene were cloned into pDM4, a Cm^r^ Ori6RK suicide plasmid (14). Primers used were 5’ GATCGTCGACATCAAATTGAATGCACTATGATC 3’ and 5’ GATCACTAGTAAAGAAGACCCCTTTATTGATTC 3’ for amplification of 1kb upstream region, and 5’ GATCACTAGTCTAGCGAGCACATAAAAAGC 3’ and 5’ GATCAGATCTTCCGGGGTGGTAAATGCTT3’ for 1kb downstream region. 1kb sequences were then inserted between SalI and SpeI sites (upstream region) or SpeI and BglII (downstream region) sites of the plasmid multiple cloning site. The resulting construct was inserted into CAB2 via conjugation by S17-1 (*λpir*) *Escherichia coli*. Transconjugants were selected for on minimal marine medium (MMM) agar containing 25 μg/mL chloramphenicol. Subsequent bacterial growth on MMM agar containing 15% (w/v) allowed for counter selection and curing of *sacB*-containing pDM4. Deletion was confirmed by PCR and sequencing analysis.

### Reconstitution of CAB2Δ*vopL*

For reconstitution of CAB2Δ*vopL*, the sequence coding for *vopL* + FLAG tag was amplified using primers 5’ GATCCTGCAGATGCTTAAAATTAAACTGCCT 3’ and 5’ GATA GAATTC TTA CTTATCGTCGTCATCCTTGTAATC CGATAATTTTGCAGATAGTGC 3’ and then cloned into the pBAD/*Myc*-His vector (Invitrogen, resistance changed from ampicillin to kanamycin) between PstI and EcoRI sites. The 1kb nucleotide sequence upstream of *vopC* (*vpa1321* in RimD2210633, accession numberNC_004605.1) was used as a promoter and cloned between XhoI and PstI sites using the primers 5’ GATC CTCGAG TATTCTTAATAAGTCAGGAGG 3’ and 5’GATC CTCGAG TATTCTTAATAAGTCAGGAGG3’. The resulting construct was inserted into CAB2Δ*vopL* via triparental conjugation using *E. coli* DH5α (pRK2073). Transconjugants were selected for on MMM agar containing 250 μg/mL kanamycin. Reconstitution was confirmed by PCR. Empty pBAD plasmid (without *vopL* gene insertion) was introduced to CAB2 and CAB2Δ*vopL* strains for consistency in bacterial strain manipulation.

### Mammalian cell culture

Caco-2 cells (ATCC, Manassas, VA) were maintained in Minimal Essential Medium with Earl’s Balanced Salts (MEM/EBSS, Hyclone, Logan, UT), supplemented with 20% (v/v) fetal bovine serum (Sigma-Aldrich, St. Louis, MO), 1% (v/v) penicillin-streptomycin (Thermo Fisher Scientific, Walthman, MA), and kept at 5% CO_2_ and 37°C. COS^phox^ cells were grown in low-glucose Dulbecco’s Modified Eagle’s Medium (DMEM, Thermo Fisher Scientific) supplemented with 10% (v/v) fetal bovine serum, 1% (v/v) penicillin-streptomycin, 1% (v/v) sodium pyruvate (Thermo Fisher Scientific), 0.8 mg/mL G418 (Thermo Fisher Scientific), 200 μg/mL hygromycin (Thermo Fisher Scientific), and 1 μg/mL puromycin (Thermo Fisher Scientific), at 37°C with 5% CO_2_ (31).

### Infection of tissue culture cells

Caco-2 and COS^phox^ cells were seeded onto 24-well plates at a density of 2.5×10^5^ (Caco-2) or 1.5×10^5^ cells/well (COS^phox^) and grown for 18-20 h. Overnight-grown bacterial cultures were normalized to an optical density at 600 nm (OD600) of 0.3 and then grown in MLB supplemented with 0.05% (w/v) bile salts for 90 min at 37°C. Growth in the presence of bile salts allowed for induction of T3SS2 (11, 12). Mammalian host cells were subsequently infected with CAB2 strains at a multiplicity of infection (MOI) of 10 using culture medium devoid of antibiotics (infection medium). To synchronize infection, cell plates were centrifuged at 200x *g* for 5 minutes. Infection was carried out for 2 h at 37 °C, after which cells were washed with unsupplemented MEM/EBSS or DMEM and subsequently treated with infection medium containing 100 μg/mL gentamicin for 1-6 h. At the end of each time point, host cells were washed with 1x PBS for removal of extracellular dead bacteria and lysed with 0.5% (v/v) TX-100. Cell lysates were serially diluted and plated on MMM agar for counting of colony forming units (CFU) as a measurement of intracellular bacterial survival/replication. To analyze bacterial filamentation, host cells were counted as containing filamentous bacteria when the cell predominantly contained bacteria longer than wild-type (CAB2) size bacteria. Filamentous bacteria needed to be at least twice the size the size of wild-type CAB2 to be considered filamentous.

Where indicated, samples were pre-treated with 100Nm GKT136901 (Aobious, Gloucester, MA) or added with 250 μM apocynin (APO, Sigma-Aldrich) at the remaining last 1 hour of infection or last hour of gentamicin incubation (6^th^ hour).

### Quantification of superoxide production

Caco-2 or COS^phox^ cells were seeded onto 6-well plates at 5×10^5^ cells/well and infected with CAB2 strains for 2 h as described above. Cells were trypsinized (0.25% trypsin/EDTA), centrifuged at 200x *g* for 5 min, and resuspended in Hank’s Balanced Salt Solution (HBSS, ThermoFisher) supplemented with Ca^2+^ and Mg^2+^. Superoxide production was measured as a function of emission of luminescence using the Diogenes kit (National Diagnostics Lab, Atlanta, GA) and according to the manufacturer’s protocol (31). Luminescence was monitored over 60 minutes using a FluoStar Optima plate reader.

### Transient transfection of COS^phox^ cells

COS^phox^ cells were transiently transfected with 0.3 μg of either wild type (WT) VopL-Flag-pSFFV or catalytically inactive VopL-WH2x3*-Flag-pSFFV constructs (16)+ 1.7 μg of empty pSFFV using Fugene HD (Promega) for 20-24 h. Subsequently, cells were treated with 0.4 μg/mL phorbol 12-myristate 13-acetate (PMA, Sigma-Aldrich) for 10 min at 37°C.

### Confocal microscopy imaging

For imaging, Caco-2 and COS^phox^ cells were seeded respectively at 2.5×10^5^ and 1-2×10^5^ cells/well onto 6-well plates containing UV-sterilized, poly-L-lysine-coated (Sigma), glass coverslips. Following the infection and transfection protocols described above, samples were fixed with 3.2% (v/v) *ρ*-formaldehyde (Thermo Fisher Scientific) for 10 minutes at room temperature. Transfected COS^phox^ cells were permeabilized with 0.5% (w/v) saponin (Sigma) for 10 minutes at room temperature and then blocked with 1% (w/v) bovine serum albumin (BSA, Sigma-Aldrich) in the presence of 0.1% saponin for 30 minutes at room temperature. In order to detect cells transfected with VopL, samples were subsequently incubated with anti-Flag antibody (1:100 dilution in 0.5% BSA, 0.1% saponin [Cell Signaling, #2368, Danvers, MA]) for 1 hour at room temperature, followed by incubation with anti-rabbit Alexa Fluor 488 conjugated secondary antibody (1:500 dilution in 0.5% BSA, 0.1% saponin [Thermo Fisher Scientific, A-21441]) for another 1h, room temperature. For detection of p67^phox^, samples were incubated with anti-p67^phox^ at 1:50 dilution (Santa Cruz, sc-7662, Dallas, TX), followed by incubation with anti-goat Alexa Fluor 555 conjugated secondary antibody (1:500 dilution [Thermo Fisher Scientific, A-21432]).

F-actin was stained with 2 units/mL of either rhodamine- or Alexa Fluor 680-phalloidin (Thermo Fisher Scientific) and DNA was stained with 1 μg/mL Hoechst A33342 (Invitrogen, Carlsbad, CA). Coverslips were placed sample-side down on glass slides containing Prolong Gold anti-fade mounting media (Thermo Fisher Scientific) and imaged on Zeiss LSM710 and LSM800 confocal microscopes. Images were converted using ImageJ (NIH).

### Statistical analysis

All data are given as mean ± standard deviation from at least 3 independent experiments unless stated otherwise. Each experiment was conducted in triplicate. Statistical analyses were performed by using unpaired, two-tailed Student’s t test with Welch’s correction. A *p* value of < 0.05 was considered significant.

## Acknowledgments

We thank Dr. Mary C. Dinauer for the valuable contribution of the COS^phox^ cell line. We also thank the members of the Orth lab for discussion and technical assistance. This work was funded by the National Institutes of Health (NIH) grant R01-AI056404, the Welch Foundation grant I-1561, and Once Upon a Time…Foundation. M.S.S. was supported by NIH grant 5T32DK007745-17. K.O. is a Burroughs Welcome Investigator in Pathogenesis of Infectious Disease, a Beckman Young Investigator, and a W. W. Caruth, Jr., Biomedical Scholar and has an Earl A. Forsythe Chair in Biomedical Science.

### Competing interests

The authors declare no conflict of interest.

